# Intertemporal choice across short and long time horizons: an fMRI study

**DOI:** 10.1101/2025.07.07.663570

**Authors:** Shengjie Xu, Jeffrey C. Erlich, Evgeniya Lukinova

## Abstract

This preregistered fMRI study investigates the neural mechanisms underlying intertemporal choices involving waiting and postponing. On a behavioral level, choices made regarding rewards available in seconds that require waiting compared to choices about rewards postponed to a number days are surprisingly similar. The explanation to this short/long time scales gap is lacking even after considering time perception and external factors, such as stress. To address that this study is the first to examine the overlapping and distinct neural circuitry involved in the intertemporal choices over seconds and days, and the waiting period within subject. Our results revealed considerable overlap in brain activation during choices that consider seconds and days delays to reward in the executive control (dACC, dlPFC) and prospection (PCC/precuneus, dmPFC) networks, but not in the valuation network. Consistent with existent literature we found the valuation network activation (both vmPFC and ventral striatum) being parametrically modulated by individual subjective values of delayed rewards. Overall, the key network determined through representational similarity and decoding analyses was prospection accounting for similarity in activation during decision making across time scales of delays and discriminating between waiting and postponing the reward. These findings enhance our understanding of the neural underpinnings of intertemporal choices and their implications for real-life decisions occurring across varying time horizons, such as paying to skip advertisements while watching videos or deciding on the next-day delivery service.

## 1 Introduction

Individual’s intertemporal preferences in the lab are predictive of important real life outcomes (Golsteyn et al., 2014; Koban et al., 2023; Mischel et al., 1989; Watts et al., 2018), which has inspired economists, psychologists and neuroscientists to investigate the behavioral and biological bases of discounting in humans and non-human animal models. However, there are two key differences in methods across species which create a gap between human and non-human studies: (i) verbal vs. nonverbal stimuli and (ii) long (days, months) vs. short (minutes, seconds) delays. Recent research established that there is a high reliability in delay discounting across the verbal/nonverbal gap, but across drastically different time scales (seconds vs. days) humans are discounting as if taking the temporal context in account: e.g., the same individual who decides to postpone twice the reward for 15 days declines to wait 30 seconds for same rewards (Lukinova & Erlich, 2021, 2024; Lukinova et al., 2019).

This fMRI study is designed to further explore the long/short time scales gap. There are numerous long delay incentive-compatible fMRI studies (Ballard & Knutson, 2009; Chen et al., 2019; Kable & Glimcher, 2007, 2010; Koban et al., 2023; Massar et al., 2015; McClure et al., 2007; McClure et al., 2004; van den Bos et al., 2014) as well as short delay experiential ones (Gregorios-Pippas et al., 2009; Jimura et al., 2013; Onoda et al., 2011; Prevost et al., 2010; Tanaka et al., 2014; Wittmann et al., 2010), some of which examined neural correlates of the waiting epoch in the short delay task. Nevertheless, to our knowledge there are no fMRI studies that investigate time preferences within subject across time scales. Our within-subject design gives us an advantage: contrasting a short delay task block to a long delay task block within runs of individual subjects could reveal differences between choices that involve waiting vs. postponing (Paglieri, 2013).

The main goal of this study is to reveal the overlapping and distinct neural circuitry underlying intertemporal choices that involve waiting for a reward vs. postponing a reward. Here, we use ‘waiting’ to refer to decisions where the delays are short and experienced and ‘postponing’ to refer to choices where rewards are put off until a future time, but other activities fill the delay duration. An example of a decision involving waiting in day-to-day life is choosing whether or not to pay to skip an advertisement before watching the video you want to watch, while one that involves postponing can be a choice whether to pay for the next-day delivery from an online shop or wait a week for free delivery.

At least three neural systems play a role in delay discounting: valuation, prospection and executive control. To date, most attention in fMRI studies on delay discounting has been focused on the valuation network (Lempert et al., 2018), in particular, Kable and Glimcher (2007) show that “neural activity in several brain regions - particularly the ventral striatum, medial prefrontal cortex and posterior cingulate cortex - tracks the revealed subjective value of delayed monetary rewards”. We expect to replicate this finding, but also to see differential interactions between valuation, prospection and executive control in the short delay vs. long delay tasks.

Our main hypotheses are as follows:

1. Overlapping brain activation will be found both in short and long temporal contexts, especially in the valuation network (ventromedial prefrontal cortex, ventral striatum and posterior cingulate cortex).
2. Less overlap between waiting and postponing in valuation, executive control and prospection networks will be observed for those subjects whose discount factor in short delay and long delay tasks are furthest from the line of best fit of log(*k_S_*) vs. log(*k_L_*) across subjects, e.g., a subject who is patient in the short but not in the long delay task will have less overlap than a subject who is patient in both tasks.
3. We expect to replicate the commonly found positive parametric modulation of the valuation network by the delayed reward, by the inverse delay of the delayed reward and by the subjective value at the time of choice (as in Kable & Glimcher, 2007).
4. On the individual level, we strongly expect to replicate the finding from Ballard and Knutson (2009) that “more impulsive subjects show less neural sensitivity to the larger magnitudes of future rewards but greater (negative) neural sensitivity to the longer delays of future rewards.”

In addition to our main hypotheses, we have several weaker predictions derived from findings in the literature that have been less consistently replicated. We anticipate:

- A.1. Negative parametric modulation of dorsolateral prefrontal cortex (executive control network) by delay during choice (at least for long delays, as in Ballard & Knutson, 2009).
- A.2. Positive parametric modulation of anterior prefrontal cortex (prospection network) by the delay during waiting (reflecting the delay-period dynamics in Jimura et al., 2013).
- A.3. Widespread activation in regions involved in executive control network (dorsolateral prefrontal cortex and dorsal anterior cingulate cortex) during waiting compared to postponing, stronger in short delay task than in the long delay task (since it will engage working memory more, as in van den Bos et al., 2014).
- A.4. Activations in prospection network (medial temporal lobe, precuneus, and dorsomedial prefrontal cortex) during waiting compared to postponing, with the signal being stronger in the long delay task than in short delay task (since prospection reduces discounting, as in Palombo et al., 2015; Peters & Büchel, 2010).

## 2 Method

This is an fMRI study pre-registered at OSF (https://osf.io/7fydc/). This study was registered prior to creation of data.

### 2.1 Participants

In this study, we recruited 97 participants for the computer session (67 female, *M_age_* = 19.67, *SD_age_* = 1.85) out of which 40 participants (27 female, *M_age_* = 19.26, *SD_age_* = 1.68; data for 31 of those (22 female) passed fMRI quality checks) were invited for the fMRI session. The selection of participants for the fMRI session was driven by: 1) fMRI safety considerations and 2) the goal to cover diverse time preferences given limited resources for the fMRI session. The latter was achieved by estimating subjects’ time preferences and choosing a subset which span a range of discount factors, but not including extremes (details in Design).

All participants were recruited from the student population of New York University (NYU) Shanghai and East China Normal University (ECNU). Subjects were informed of the opportunity to participate through recruitment flyers posted on the university website, campus bulletin boards, and WeChat. Subjects that were willing to participate in the study scanned the QR code, read the brief intro, and filled in the registration form. All forms, instruction, and stimuli were in Chinese.

The recruitment and experimental sessions for the study were completed between 6th October 2019 and 26th March 2021 (impacted by COVID-19). The study was approved by the Institutional Review Board (IRB, protocol 024-2017) of NYU Shanghai. All subjects gave informed consent (written) before participation in the study. All research was performed in accordance with relevant guidelines and regulations.

### 2.2 Sample Size

The target sample size (for the fMRI session) was 40 participants. Standard fMRI study includes around 30 participants, e.g., 32 (as in Tanaka et al., 2014) after discarding subjects whose data contains too many artifacts. No power analysis was used to set the sample size. The target sample size was set based on the funding constraints. We estimated that our effect size would be comparable to previous within-subject studies of intertemporal choice (Lamichhane et al., 2022; Peters & Büchel, 2010).

### 2.3 Procedure

The participation in the computer session took from 1 hour to 1.5 hours, including a short survey collecting demographic information and placing participants on the Barratt Impulsiveness Scale (BIS-11; Patton, Stanford, et al., 1995). The total time for participation in the fMRI session was approximately 1.5 hours, including ∼ 15 minutes for filling out the forms. Subjects were compensated for 20 RMB for any level of participation in the computer session and for 100 RMB in the fMRI session. In addition, they could receive from 0 RMB up to 50 RMB through the reward system tied to the task.

### 2.4 Design

First, all subjects participated in the computer session and then they might or might not be invited to the fMRI session on a separate day. The tasks and the stimuli appearance used across the sessions were the same. All subjects watched a refresher video before the start of the fMRI session. In the intertemporal choice task, two options were presented on the screen in each trial: a delayed option and an immediate option (Figure 1). Subjects made a choice given reward magnitude of both options (*reward*, the immediate option *reward* was 4 coins for all trials) and delay (*delay*, in seconds or in days) of the delayed option. ‘Short Delay Task’ (Short, S) had delays in seconds and ‘Long Delay Task’ (Long, L) had delays in days.

**Figure 1:**
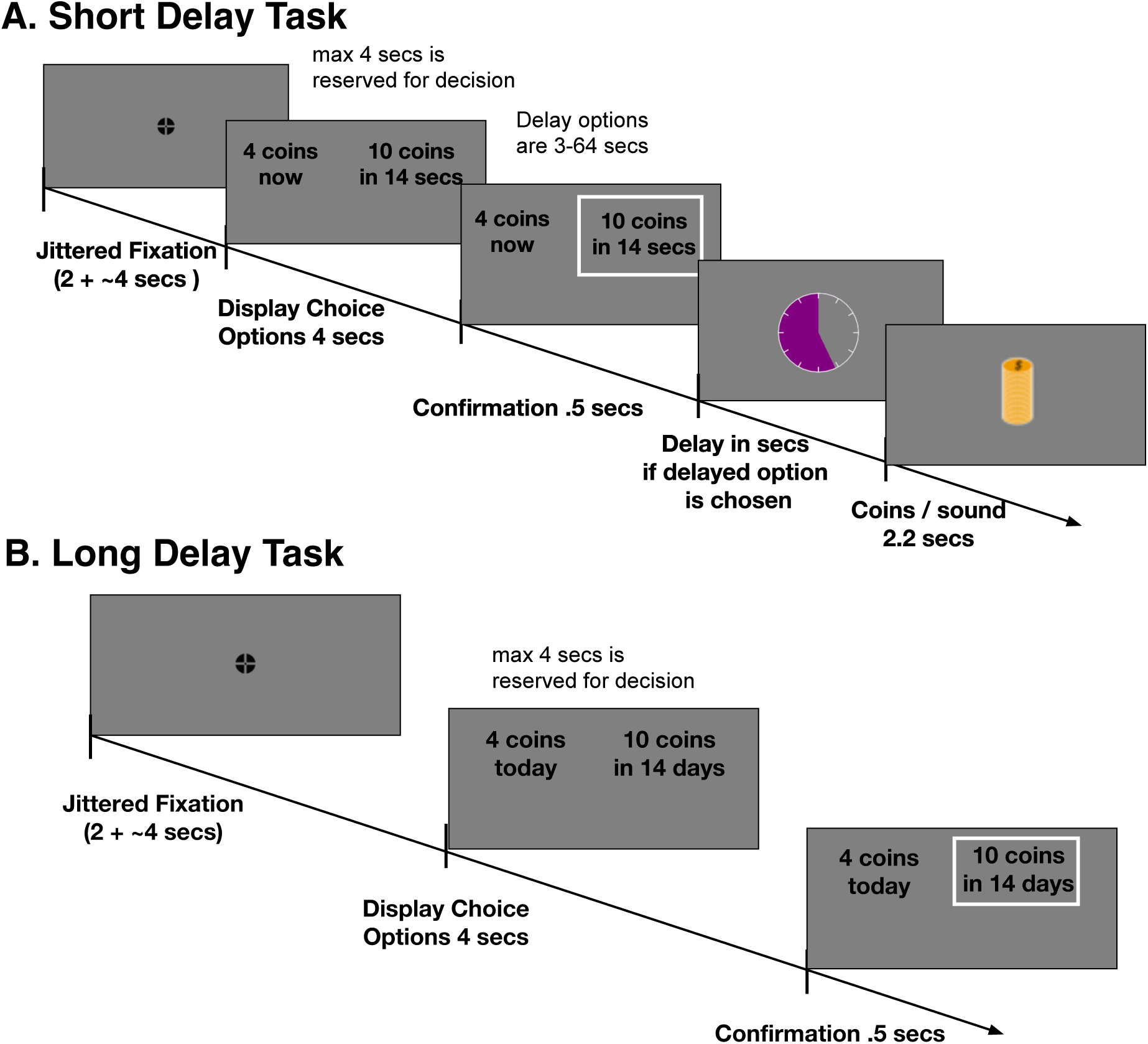
Timelines of the experimental tasks. **A. Short Delay Task.** The total timing of the trial was dependent on the delay if delayed option was selected: 2 + 4 + 4 + .5 + 2.2+ **delay** ≈ (12.7+ **delay**) seconds per trial. **B. Long Delay Task.** The total timing of the trial was independent of selected options: 2 + 4 + 4 + .5 ≈ 10.5 seconds per trial.

The behavioral criteria we used to determine whether to invite subjects to the fMRI session or not were the following: in the computer session (i) subjects made at least 10% choices of ‘immediate’ and ‘delayed’ category in both Short and Long tasks and (ii) subjects were sensitive to reward and delay. We obtained *choice* (a binary variable, where 1 is for delayed option and 0 for immediate option), *reward*, and *delay* variables per trial of the task and first checked whether for each task 0.1 ≤ *M* (*choice*) ≤ 0.9. Then we also examined whether both *p*-values (for *reward* and *delay* coefficients) in the regression (*choice* ∼ *reward* + *delay*) for each task were significant. In other words, subjects were excluded if they almost always chose the delayed (or immediate) option or if they were insensitive to either delay or reward.

All participants in our within-subject study did exactly the same tasks (in a different order, described later). We used a mixed design type, i.e. a block design with events within the blocks. In the computer session, subjects did 100 trials of each task (200 trials total) divided into 2 runs (2 blocks(tasks) per run) of 50 trials of Short and 50 trials of Long each in an interleaved fashion: S-L-S-L or L-S-L-S for half of the subjects similar to our previous studies (Lukinova et al., 2019). In the fMRI session, subjects also completed 100 trials of each task (200 trials total), divided into 4 runs (2 blocks/tasks per run) of 25 trials of Short and 25 trials of Long each in an interleaved fashion: S-L-L-S-S-L-L-S or L-S-S-L-L-S-S-L. Therefore, we acquired a total of 200 trials per subject per session. The total length of the Short Delay Task was directly dependent on subject’s choices, i.e. whether they decided to (actively) wait or not. The choice sets (reward magnitudes and delays) were optimized to be able to fit the diverse time preferences of subjects (the simulation is described in the preregistration). The payment for the experiment was incentive-compatible: subjects were paid according to their real choices (in S all trials were counted for additional payment, 1 coin = 0.02 RMB; in L one trial was chosen randomly for payment, 1 coin = 2 RMB).

In the Short Delay Task (Figure 1A) the trial started with the jittered fixation (event ‘fixation’), followed by the choice options being displayed for a maximum of 4 seconds (event ‘decision’), then the confirmation of the choice was displayed (event ‘confirmation’), finally either the coins sound and the visualization (event ‘coins’) was presented right away (if the immediate option was chosen) or after a certain delay in seconds (if the delayed option was chosen, event ‘waiting’ with a clock appearing on the screen). In the Long Delay Task (Figure 1B) the trial also started with the jittered fixation (event ‘fixation’), followed by the choice options presented for a maximum of 4 seconds (event ‘decision’), and the confirmation (event ‘confirmation’). No coins or clock were shown after these events since only one random trial counted for additional payment, which was determined at the end of the session. The timeline and the expected trial duration of the Short and Long tasks are shown in Figure 1A and Figure 1B, respectively. The responses for fMRI session were collected using a button box placed in participant’s right hand, where left choice was attached to button #1 (index finger) and right choice was attached to button #2 (middle finger) (computer session - keys ‘left arrow’ and ‘right arrow’ on the keyboard, respectively).

### 2.5 Behavioral Analysis

Following our established procedures (Lukinova et al., 2019), we fitted subjects’ choices using a Bayesian hierarchical model (BHM) of hyperbolic discounting with decision noise. The model had four population-level parameters (log discount factor, log(*k*), and decision noise, log(*τ*), for each of the two tasks, also known as fixed effects) and three parameters per subject: log(*k_S_*), log(*k_L_*), and log(*τ*).

choice ∼ (rewmag /(1 + exp (logk) * delay) - smag)/ exp (noise),

noise ∼ unit + (1 | subjid),

logk ∼ unit + (unit | subjid)

where rewmag was the later *reward*; smag was the sooner *reward*; unit was the task separator by the unit of *delay*; logk was the natural logarithm of the discount factor *k*; and noise was the natural logarithm of the decision noise, or log(*τ*). The same fitting procedure was done for the computer and the fMRI sessions (separately). The range of discount factors was consistent with that reported in our prior research (Figure 2A).

**Figure 2:**
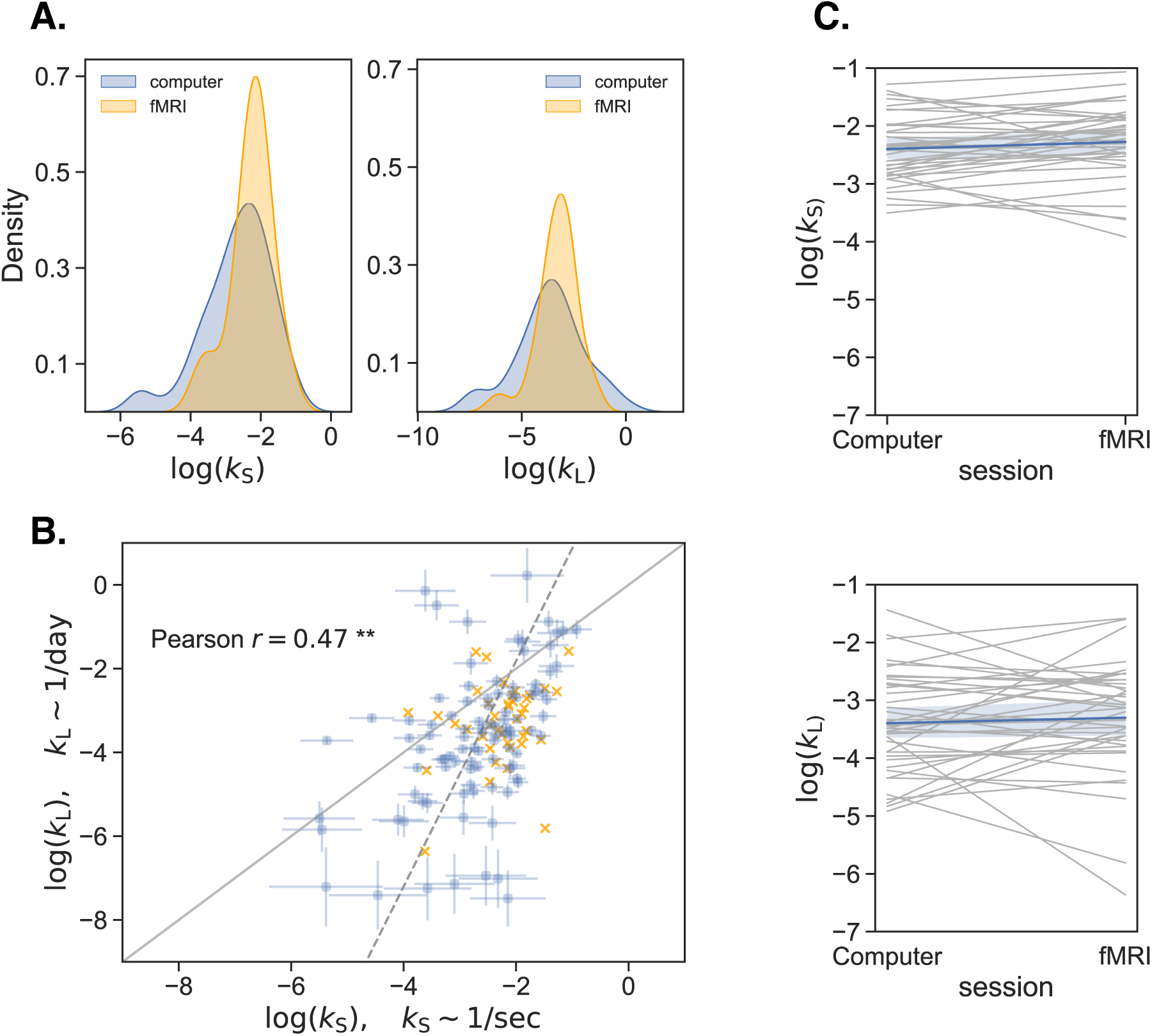
Behavioral results. **A.** Distribution of discount factors between computer and fMRI sessions across Short and Long tasks. **B.** Time preferences across temporal contexts. Each circle was one subject in the computer session (*N* = 97). The error bars were the SD of the estimated coefficients. The lines shown included the unity line (*y* = *x*) and the perpendicular least squares regression line. The logs of discount factors (log(*k*)) in S (x-axis) were plotted against the ones in L (y-axis). The Pearson correlation reported on the plot was for the computer session (*r* = 0.47, ** for *p <* 0.001). Each ‘*X*’ represented the discount factors of each subject in the fMRI session (*N* = 40). **C.** Stability of discount factors across sessions in Short (top) and Long (bottom). Each subject’s log(*k*) (y-axis) was plotted via a slopegraph (with x-axis being an experimental session). The mean and 95% confidence interval were plotted using color.

After eliciting discount factors, subjective value (*SV*) was calculated for the delayed option per trial, per task, per subject, where individual subjective value of the delayed reward was assumed to be hyperbolic 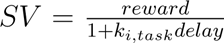, and *k_i,task_* represented the discount factor for the *i^th^*subject in either the Short or the Long *task*.

The permutation tests of within-subject differences between the computer and the fMRI sessions were done by shuffling the group label and computing the mean between the shuffled groups 10000 times. This generated a null distribution which was used to estimate the probability of observing the true difference from 0. Whenever multiple hypotheses were tested at the same time (e.g., multiple correlations were run between model-based and model-free estimates) we used the Bonferroni multiple comparison correction with *α*-level being *α/n*, where *n* was the number of hypotheses.

### 2.6 fMRI Analysis

#### 2.6.1 fMRI Acquisition

At least two research assistants interacted with subjects during the fMRI session. First, the localizer and the structural scan sequences were run, in this order. Then, fMRI scanning was divided into four functional runs. The fMRI technician was asked to set the scan time to 20 mins, but the actual duration of the scan differed depending on the subject’s choice to wait or not. The technician was asked to stop scanning when one run of the behavioral task was over. After two runs, subject could take some rest, if necessary.

Functional MR images (whole brain) were obtained for each subject by using Siemens Prisma 3T system at the Neuroimaging Center of East China Normal University (ECNU). Images were acquired by using echo-planar T2 images (EPI, gradient echo (GRE), using Simultaneous Multi-Slice (SMS) package: GRAPPA 2, MB 2, No PF) with BOLD (blood oxygenation-level-dependent) contrast, and angled 30^◦^ with respect to the AC-PC line to minimize susceptibility artifacts in the orbitofrontal cortex (Deichmann et al., 2003). MR imaging settings were as follows: repetition time (TR) = 1,740 ms; echo time (TE) = 27.0 ms; slice thickness = 3 mm yielding a 64 x 64 x 42 matrix (3 mm x 3 mm x 3 mm); flip angle = 90^◦^; FOV read = 192 mm; FOV phase = 100 %, interleaved series order (the exact sequence parameters are in the attached file). The EPI sequence was optimized to reduce the dropoff in OFC area. High-resolution structural T1-weighted scans (0.9 mm x 0.9 mm x 0.9 mm) were acquired by using an MPRage sequence. Visual stimuli were presented by means of a mirror mounted on the MRI head coil, and responses were acquired via an MRI-safe 5-button response box (Sinorad).

The MRI data collected was in DICOM format that was converted into 4d NIfTI format (BIDS) by using ‘dcm2nii’ (through ‘MRIcroGL’). The data then underwent a QC monitoring (described in detail in the next section). Next, we used ‘3dDespike’ (default options) from AFNI (Analysis of Functional NeuroImages, version 19.1.08 ‘Caligula’ Cox (1996)) to remove ‘spikes’ from the data that had fd num*>*0 (from QC), i.e. data that had some spikes due to motion. This software “writes a new dataset with the spike values replaced by something more pleasing to the eye” and stores the data in the AFNI data format (BRIK+HEAD). To convert back to the BIDS format for preprocessing we used ‘3dAFNItoNIFTI’ (AFNI).

#### 2.6.2 fMRI Quality Check

Before running preprocessing, several quality checks were implemented to ensure that results were not influenced by artifacts. We performed quality control of MRI data in two ways: i) a quantitative check with ‘mriqc’ package (version 0.15.0) at the group level to have both individual metrics and group-level analysis to find outliers across all subjects (Esteban et al., 2017) and ii) a visual qualitative check (details in the preregistration). If a BOLD run did not pass this check, this BOLD run was removed from further analysis. If more than two runs were excluded for a single subject, that subject was excluded. If a structural T1-weighted scan did not pass this check, that subject was excluded. Data for 31 participants was left at this stage.

#### 2.6.3 fMRI Preprocessing

*The text in this subsection was automatically generated by fMRIPrep:* Results included in this manuscript come from preprocessing performed using *fMRIPrep* 23.2.3 (RRID:SCR 016216; Esteban et al., 2018, 2019), which is based on *Nipype* 1.8.6 (RRID:SCR 002502; K. Gorgolewski et al., 2011; K. J. Gorgolewski et al., 2018).

##### Anatomical data preprocessing

A total of 1 T1-weighted (T1w) images were found per subject within the input BIDS dataset. The T1w image was corrected for intensity non-uniformity (INU) with N4BiasFieldCorrection (Tustison et al., 2010), distributed with ANTs 2.5.0 (Avants et al., 2008, RRID:SCR 004757), and used as T1w-reference throughout the workflow. The T1w-reference was then skull-stripped with a *Nipype* implementation of the antsBrainExtraction.sh workflow (from ANTs), using OASIS30ANTs as target template. Brain tissue segmentation of cerebrospinal fluid (CSF), white-matter (WM) and gray-matter (GM) was performed on the brain-extracted T1w using fast (FSL (version unknown), RRID:SCR 002823, Y. Zhang et al., 2001). Brain surfaces were reconstructed using recon-all (FreeSurfer 7.3.2, RRID:SCR 001847, Dale et al., 1999), and the brain mask estimated previously was refined with a custom variation of the method to reconcile ANTs-derived and FreeSurfer-derived segmentations of the cortical gray-matter of Mindboggle (RRID:SCR 002438, Klein et al., 2017). Volume-based spatial normalization to one standard space (MNI152NLin2009cAsym) was performed through nonlinear registration with antsRegistration (ANTs 2.5.0), using brain-extracted versions of both T1w reference and the T1w template. The following template was selected for spatial normalization and accessed with *TemplateFlow* (23.1.0, Ciric et al., 2022): *ICBM 152 Nonlinear Asymmetrical template version 2009c* [Fonov et al. (2009), RRID:SCR 008796; TemplateFlow ID: MNI152NLin2009cAsym].

##### Functional data preprocessing

For each of the 4 BOLD runs found per subject (across all tasks and sessions), the following preprocessing was performed. First, a reference volume was generated, using a custom methodology of *fMRIPrep*, for use in head motion correction. Head-motion parameters with respect to the BOLD reference (transformation matrices, and six corresponding rotation and translation parameters) were estimated before any spatiotemporal filtering using mcflirt (FSL, Jenkinson et al., 2002). The BOLD reference was then co-registered to the T1w reference using bbregister (FreeSurfer) which implements boundary-based registration (Greve & Fischl, 2009). Co-registration was configured with six degrees of freedom. Several confounding time-series were calculated based on the *preprocessed BOLD*: framewise displacement (FD), DVARS and three region-wise global signals. FD was computed using two formulations following Power (absolute sum of relative motions, Power et al. (2014)) and Jenkinson (relative root mean square displacement between affines, Jenkinson et al. (2002)). FD and DVARS were calculated for each functional run, both using their implementations in *Nipype* (following the definitions by Power et al., 2014). The three global signals were extracted within the CSF, the WM, and the whole-brain masks. Additionally, a set of physiological regressors were extracted to allow for component-based noise correction (*CompCor*, Behzadi et al., 2007). Principal components were estimated after high-pass filtering the *preprocessed BOLD* time-series (using a discrete cosine filter with 128s cut-off) for the two *CompCor* variants: temporal (tCompCor) and anatomical (aCompCor). tCompCor components were then calculated from the top 2% variable voxels within the brain mask. For aCompCor, three probabilistic masks (CSF, WM and combined CSF+WM) were generated in anatomical space. The implementation differed from that of Behzadi et al. in that instead of eroding the masks by 2 pixels on BOLD space, a mask of pixels that likely contained a volume fraction of GM was subtracted from the aCompCor masks. This mask was obtained by dilating a GM mask extracted from the FreeSurfer’s *aseg* segmentation, and it ensured components were not extracted from voxels containing a minimal fraction of GM. Finally, these masks were resampled into BOLD space and binarized by thresholding at 0.99 (as in the original implementation). Components were also calculated separately within the WM and CSF masks. For each CompCor decomposition, the *k* components with the largest singular values were retained, such that the retained components’ time series were sufficient to explain 50 percent of variance across the nuisance mask (CSF, WM, combined, or temporal). The remaining components were dropped from consideration. The head-motion estimates calculated in the correction step were also placed within the corresponding confounds file. The confound time series derived from head motion estimates and global signals were expanded with the inclusion of temporal derivatives and quadratic terms for each (Satterthwaite et al., 2013). Frames that exceeded a threshold of 0.5 mm FD or 1.5 standardized DVARS were annotated as motion outliers. Additional nuisance timeseries were calculated by means of principal components analysis of the signal found within a thin band (*crown*) of voxels around the edge of the brain, as proposed by (Patriat et al., 2017). All resamplings could be performed with *a single interpolation step* by composing all the pertinent transformations (i.e. head-motion transform matrices, susceptibility distortion correction when available, and co-registrations to anatomical and output spaces). Gridded (volumetric) resamplings were performed using nitransforms, configured with cubic B-spline interpolation.

Many internal operations of *fMRIPrep* use *Nilearn* 0.10.2 (Abraham et al., 2014, RRID:SCR 001362), mostly within the functional processing workflow. For more details of the pipeline, see the section corresponding to workflows in *fMRIPrep*’s documentation.

#### 2.6.4 fMRI Statistical Modeling

We used Nilearn (Abraham et al., 2014) for all statistical modeling analyses.

##### GLM Analysis

For all general linear models (GLM) we set haemodynamic response function to double-gamma SPM and a lag-1 autoregressive model - ar(1) - for the temporal structure of the noise on the individual level. On the group level we thresholded a z-statistic image with FDR *<.*05 (False Discovery Rate) and cluster size *>* 30.

In the first model (GLM1), we did a localization analysis of temporal context, i.e. we located areas that corresponded to decision epoch in the tasks, and waiting versus postponing. This model addressed the main hypothesis 1 (overlapping brain activations for both tasks in valuation network), hypothesis 2 (less overlap between short and long tasks for those subjects further away from the fitted line of log(*k_S_*) vs. log(*k_L_*)) and weaker hypotheses A.3 and A.4 (activations in executive control and prospective networks, respectively). We modeled decision (from the onset of the event till confirmation) and confirmation (from the onset of confirmation event until the onset of the new trial) epochs in a corresponding trial, resulting in two regressors in each task.

Therefore, the design matrix contained four regressors of interest (Figure 3A). Six motion regressors, FD, white matter, cerebrospinal fluid (CSF) signal, and cosine values for each TR - generated by fMRIPrep during the preprocessing step - were included as regressors of no interest. Eight linear contrasts are displayed in Table 1. Contrasts 1-4 on the group level generated z-statistic images for each of the regressors of interest. Contrasts 5, 7 (6, 8) showed where in the brain activations are stronger for Short (Long). On the group level one-sample t-test was performed for each contrast (i.e. does the group activate on average?) addressing the main hypothesis 1. Overlap was calculated based on shared thresholded clusters in Short and in Long (ranked by peak statistic in both tasks). Considering two groups (one, *N* = 15, closest to the fitted line of log(*k_S_*) vs. log(*k_L_*) and the other, *N* = 16, - furthest) we performed a two-sample t-test for contrasts 5-8 (i.e. is there a significant group difference?) that accounted for the main hypothesis 2.

**Figure 3:**
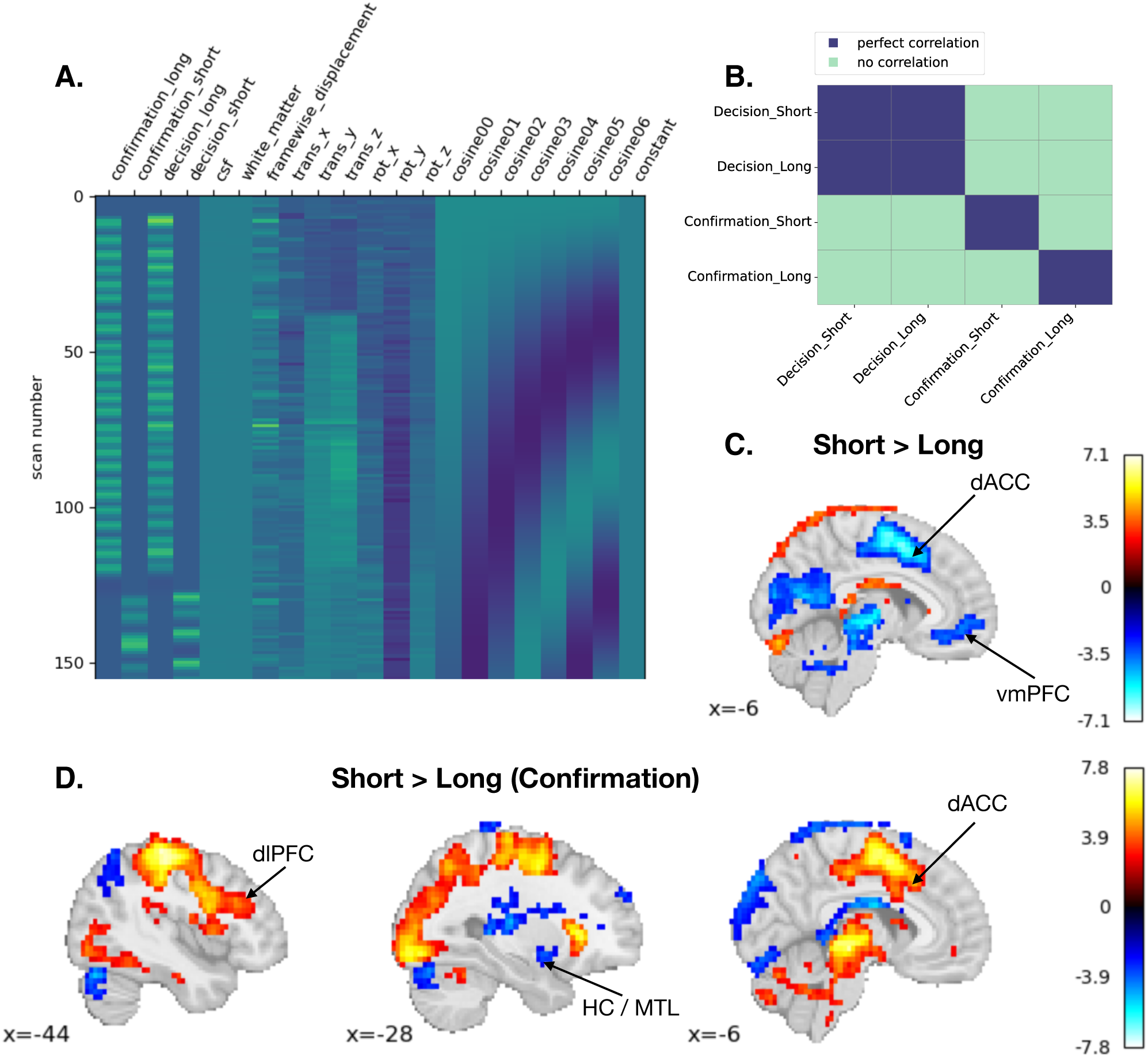
Imaging data modeling and GLM1 results. **A.** GLM1 design matrix for an example subject (cut just after 150 scans). First four columns were regressors of interest. **B.** Theoretical matrix for representational similarity analysis (RSA) that accounted for similarity between decision epoch across tasks but differentiated between confirmation epochs, i.e. waiting and postponing. **C.** Thresholded z-statistic map of contrast 5 (Decision, Short *>* Long) at *x* = −6. Activations in dACC (executive control) and vmPFC (valuation) were smaller in Short than in Long during decision. Colormap represented z-statistic value. **D.** Thresholded z-statistic map of contrast 7 (Confirmation, Short *>* Long) at *x* = [−44, −28, −6]. There was more engagement in dlPFC (executive control) and dACC (executive control), while less engagement in HC/MTL (prospection) when waiting was compared postponing.

**Table 1:** GLM1, GLM2 and GLM2c contrasts.

| Regressors -> | Decision Short | Decision Long | Confirmation Short | Confirmation Long |
| --- | --- | --- | --- | --- |
| GLM1 contrasts |  |  |  |  |
| 1. Decision Short | 1 | 0 | 0 | 0 |
| 2. Decision Long | 0 | 1 | 0 | 0 |
| 3. Confirmation Short | 0 | 0 | 1 | 0 |
| 4. Confirmation Long | 0 | 0 | 0 | 1 |
| 5. Decision Short > Decision Long | 1 | -1 | 0 | 0 |
| 6. Decision Long > Decision Short | -1 | 1 | 0 | 0 |
| 7. Confirmation Short > Confirmation Long | 0 | 0 | 1 | -1 |
| 8. Confirmation Long > Confirmation Short | 0 | 0 | -1 | 1 |
| GLM2 contrasts |  |  |  |  |
| 1. Decision Short | 1 | 0 |  |  |
| 2. Decision Long | 0 | 1 |  |  |
| 3. Decision Short > Decision Long | 1 | -1 |  |  |
| GLM2c contrasts |  |  |  |  |
| 1. Confirmation Short |  |  | 1 | 0 |
| 2. Confirmation Long |  |  | 0 | 1 |
| 3. Confirmation Short > Confirmation Long |  |  | 1 | -1 |

Then, the GLM1 was run one more time omitting the smoothing step (GLM1u). This was done to avoid blurring the signal between voxels: keeping each voxel as distinct as possible would help decoding techniques to distinguish between patterns elicited by different stimuli. We used contrasts 1-4 on the individual level from GLM1u as MVPA inputs.

Finally, we estimated a separate general linear model (GLM1s) in line with the Least-Squares Separate (LSS) approach (Mumford et al., 2012), where the trial of interest was modeled with its own regressor and all other trials were grouped into a second regressor. This trial-wise modeling approach was used to generate beta maps that more accurately capture trial-specific neural activity and served as input to subsequent decoding analysis.

The parametric modulation modeling (GLM2-4), i.e. assuming BOLD response amplitude was dependent on certain parameters’ values, was implemented in Nilearn via a ‘modulation’ column (equivalent to including unmodulated regressor per one de-meaned modulated one). These models tested the main hypothesis 3.

In GLM2, we modeled one regressor (from the decision onset till the confirmation) for each task as slopes with de-meaned reward magnitudes. The same regressor was used for confirmation (from the onset of confirmation until the onset of the new trial) in a separate model, GLM2c. In GLM3 (GLM3c), we considered parametric modulation by delay, whereas in GLM4 parametric modulation by subjective value (*SV*). The design matrix resembled the GLM2 but with a different de-meaned modulated regressor. The event modeled was incorporated in the model’s name (i.e. ‘…c’ for confirmation). Contrasts were similar for all models above, e.g., GLM2(c) in Table 1.

##### ROI Analysis

The ROI analysis focused on the valuation network. We used the conjunction mask from Hare et al. (2014). This mask included voxels in ventromedial prefrontal cortex (vmPFC), ventral striatum, and posterior cingulate cortex (PCC). In particular, we addressed hypothesis 1 by extracting individual beta-maps for this ROI for contrasts 1 and 2 from the GLM1. Then, for each participant, we ran a correlation across all voxels in this region to see how similar the pattern of activation was during Short and Long tasks. One participant was excluded from this analysis with all voxels having zero *β* coefficients.

From GLM2(c), GLM3(c) and GLM4, we extracted individual beta-maps of contrasts 1 and 2 from the peak voxels (determined via thresholded z-maps) within the ROI and computed average values of the modulated parameter estimates for reward, delay, and *SV*, respectively, separately for decision and confirmation epochs, and across tasks. Then we 1) ran a one-sample t-test (comparing *β*-coefficient averages to 0) to address hypothesis 1; and 2) ran a two-sample t-test to compare the two groups (closest to the fitted line of log(*k_S_*) vs. log(*k_L_*) and furthest) to address hypothesis 2.

##### Individual Differences Analysis

Using parameter estimates from GLM2, GLM3 and GLM4, we examined whether individuals’ neural responsiveness to reward magnitude, delay, and *SV* correlated with their discount factor log(*k*). We extracted individual beta-maps of contrasts 1 and 2 from each parametric model within the valuation network separately and averaged them across voxels. We then correlated these individual-averaged *β* coefficients against the individuals’ tendency to discount the future option, log(*k*), (as in Figure 4: Ballard & Knutson, 2009) to test the main hypothesis 4.

**Figure 4:**
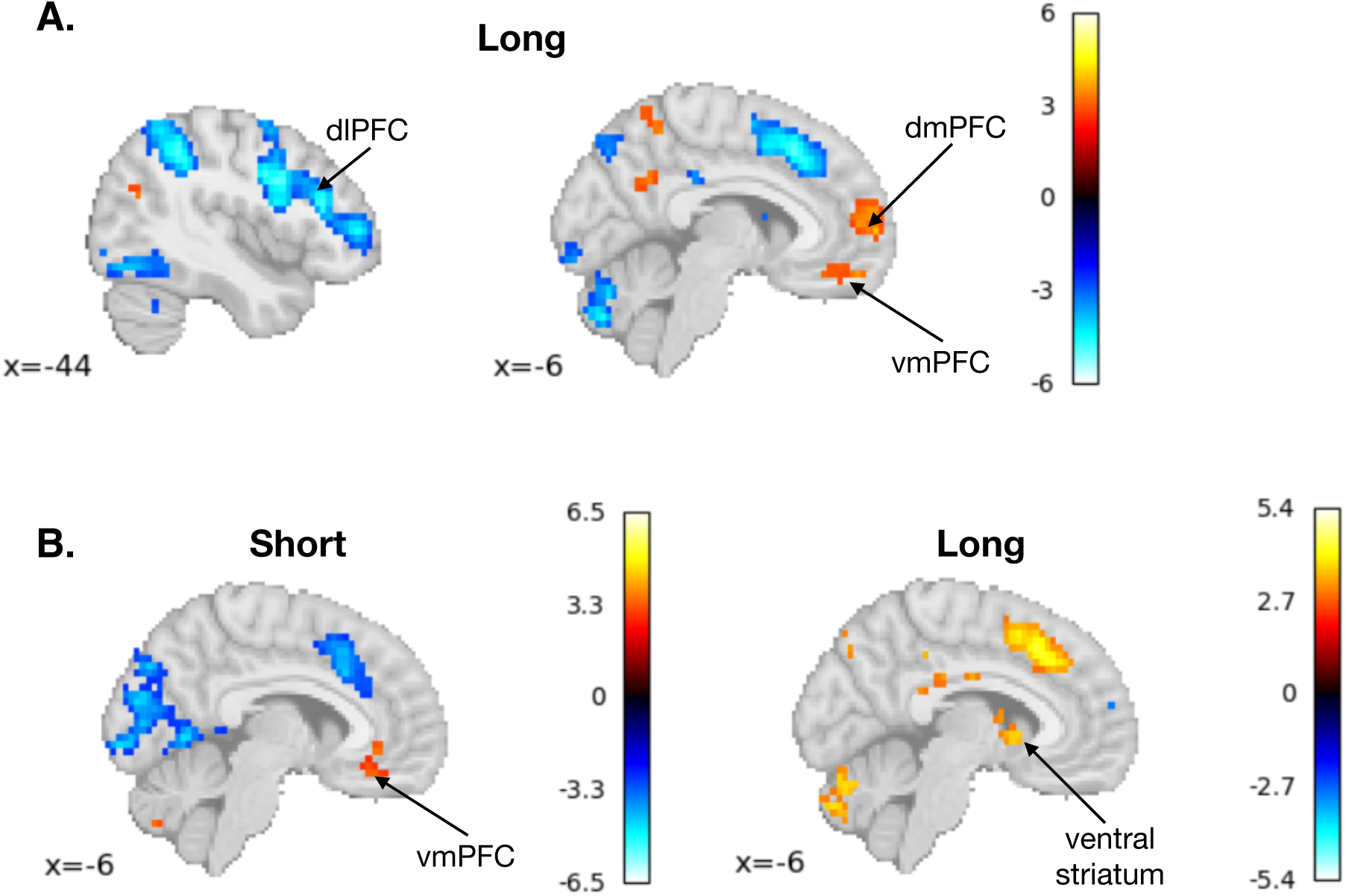
Parametric modeling results. **A.** Neural sensitivity to delay in the Long task. Thresholded z-statistic map for GLM3 at *x* = [−44, −6]. Similar to Short, in the Long task we found negative modulation - the bigger the delay magnitude the smaller the activation - in dlPFC (executive control) by delay magnitude and positive modulation in dmPFC (prospection) and vmPFC (valuation) during decision making. Colormap represented z-statistic value. **B.** Neural sensitivity to subjective value (*SV*). Thresholded z-statistic map for Short (left) and Long (right) at *x* = −6. Both ventral striatum and vmPFC were parametrically modulated by *SV* on the individual level across tasks, with prevalence of activations defining which region would win on the group level.

#### 2.6.5 Multi-voxel Pattern Analysis

With multi-voxel pattern analysis (MVPA), we examined to what extent Short and Long tasks evoke similar neural patterns.

##### RSA

We performed representational similarity analysis (RSA; RSA toolbox, Nili et al., 2014) to examine the encoding of waiting, postponing, Short and Long tasks using multi-voxel patterns. To compute dissimilarity, we used Pearson correlational distance, which is insensitive to changes of overall activation magnitude, therefore, makes the RSA orthogonal to the univariate GLM analysis (Walther et al., 2016). As a part of our planned analysis, we used this methodology combined with a whole-brain “searchlight” procedure (Kriegeskorte et al., 2006; Z. Zhang et al., 2017), in which we examined patterns in the immediate neighborhood of every voxel (a 27-voxel cube with that voxel as the center) in the brain. This procedure allowed us to find where in the brain certain task-related variables were encoded, except that now we focused on the multi-voxel pattern of activation, rather than the activation magnitude in single voxels after GLM1u. We extracted beta-maps for contrasts 1-4 from the GLM1u individual-level results. Therefore, the activation patterns of a 27-voxel searchlight for all 4 contrasts of interest could be represented by four 27-dimensional vectors of *β* coefficients. For each searchlight, we generated a 4 x 4 neural representational dissimilarity matrix (neural RDM) for each individual participant, based on the Pearson correlational distance between activation patterns for all possible pairs of conditions. We compared the neural RDMs to hypothesized dissimilarities in neural patterns between conditions based on a theoretical matrix (Figure 3B) that accounted for similarity between decision epoch of Short and Long tasks but differentiated between confirmation epochs, i.e. waiting and postponing, via the whitened cosine similarity (Diedrichsen et al., 2020) to address hypotheses 1 and 2. We assessed statistical significance using a permutation-based approach (1000 permutations). The p-value was computed as the fraction of permutations in which the shuffled neural RDMs yielded a whitened cosine similarity greater than or equal to the observed value. The p-values were then Fisher-transformed to z-values and thresholded with FDR *<* .05 and the cluster size *>* 30.

##### Decoding

In our preregistered analysis, we planned to check whether Short and Long tasks have indeed a common code or their representations in the brain are distinct at a more fine-grained level via a cross-categorical decoding approach (as in Kobayashi & Hsu, 2019).

We used Nilearn (Abraham et al., 2014), individual beta-maps from GLM1u, and individual trial-wise beta maps from GLM1s in the decoding analyses. The decoding was performed either considering one brain network at a time or a joint mask consisting of all three networks of interest. Masks for executive control (the dorsolateral prefrontal cortex (dlPFC), the dorsal anterior cingulate cortex (dACC)) and prospection (the dorsomedial prefrontal cortex (dmPFC), PCC/precuneus) networks were created by combining respective areas via Multi-subject Dictionary learning probabilistic atlas (MSDL, Varoquaux et al., 2011). In addition, areas of hippocampus (HC) / and medial temporal lobe (MTL) were added to the prospection mask using Julich-Brain probabilistic atlas (Amunts et al., 2020).

First, in order to perform cross-categorical decoding we trained a support vector regression (SVR) to decode subject-by-subject absolute difference in log(*k*) between tasks, | log(*k_Short_*) − log(*k_Long_*)|, from voxel-level activation patterns (GLM1u) of one task (e.g., Short) using leave-one-subject-out cross-validation. Then, the best-fit model based on the mean absolute error (MAE) was used to predict | log(*k_Short_*) − log(*k_Long_*)| using the beta-maps from the other task (e.g., Long, and vice versa). The procedure was run using masks of each network of interest separately.

As preregistered, we also decoded trial-by-trial *SV* from trial-wise beta maps derived from first-level GLM1s estimates of BOLD responses in a “searchlight” of a 10-mm radius (Kriegeskorte et al., 2006) within the joint mask. We used an SVR with leave-one-run-out cross-validation per participant for Long and Short tasks separately. Then, we aggregated prediction performance by computing average voxel-wise maps of R² scores across participants.

In addition, we applied several classification models (support vector classifier (SVC), logistic regression, and ridge classifier) to check how well we can classify the tasks (Short vs Long) with beta-maps from GLM1u within valuation, executive control, and prospection network masks separately. Classification was performed using nested cross-validation, with a 5-fold inner loop for hyperparameter tuning based on classification accuracy within the model class, and a 4-fold outer loop for evaluating the same class model performance on held-out data resulting in CV-training and CV-testing accuracies.

This approach allowed us to test whether all voxels in the mask were helpful in classifying the Short and the Long tasks. Still, we aimed to know which regions were driving the classification accuracy. So, we ran a “searchlight” of a 10-mm radius within each network for the best classifier with 5-fold cross-validation to identify the key clusters within the masks.

### 2.7 Discrepancies from Preregistered Procedures

We used Nilearn (as opposed to FSL) for statistical analysis of imaging data. Parametric models were split to model decision and confirmation events separately (e.g., GLM2 and GLM2c). Decoding procedures were enhanced beyond SVR (preregistered) and regression in general. Cross-categorical decoding approach was modified, predicting | log(*k_Short_*) − log(*k_Long_*)| (as opposed to *SV*).

## 3 Results

### 3.1 Behavioral Data

We collected behavioral data via a computer session (*N* = 97) and an fMRI session (*N* = 40, Figure 1) where participants made choices in the delay-discounting task in seconds (Short) and in days (Long). Time preferences were elicited by fitting participants’ choices with a Bayesian hierarchical model (BHM) of hyperbolic discounting with decision noise (Method). The ranges of individual discount factors (log(*k*)) were similar to those reported previously and had a smaller range for the fMRI session compared to the computer one in Figure 2A (by design, Method).

Consistent with results from our prior research (Lukinova & Erlich, 2021, 2024; Lukinova et al., 2019) we found that in the computer session log(*k*) was significantly correlated between Short and Long tasks (Pearson *r* = 0.47, *p <* 0.001; Figure 2B). Significant correlations were found between model-based and model-free measurements, i.e. the log(*k*) was negatively correlated with the proportion of later-larger (*pLL*) choices within the tasks (in Short: Pearson *r* = −0.94, *p <* 0.001; and in Long: Pearson *r* = −0.95, *p <* 0.001) as well as cross-task correlations remained significant (log(*k_S_*) vs *pLL_L_*: Pearson *r* = −0.49, *p <* 0.001; log(*k_L_*) vs *pLL_S_*: Pearson *r* = −0.46, *p <* 0.001). We found no correlation between delay-discounting measurements and BIS (all *ps >* 0.390), or decision response time (RT, all *ps >* 0.260), while across tasks RTs were significantly correlated (*RT_L_*vs *RT_S_*: Pearson *r* = 0.66, *p <* 0.001).

Within the fMRI session we did not observe a significant correlation between discount factors across tasks (Pearson *r* = 0.26, *p* = 0.105), partially due to the criteria for choosing subjects for the fMRI session (Figure 2A and B, Method). Again, although RTs were significantly correlated between the tasks (*RT_L_* vs *RT_S_*: *r* = 0.84, *p <* 0.001), response times were not correlated to the discount factors (all *ps >* 0.510). For subjects who participated in the fMRI session we checked the stability of their time preferences across sessions. According to the permutation tests log(*k*) was not significantly different across the sessions (in Short: *M* (log(*k_computer_*)) = −2.40 & *M* (log(*k_fMRI_*)) = −2.27, *p* = 0.101; and in Long: *M* (log(*k_computer_*)) = −3.40 & *M* (log(*k_fMRI_*)) = −3.30, *p* = 0.491; also plotted in Figure 2C).

### 3.2 fMRI Data

We first tested which brain regions were activated for all subjects as a group during the decision epoch for Short and Long tasks. There was a considerable overlap in the areas of activation, including the areas of dlPFC, dACC, and dmPFC (Table 2). The brain activations were smaller in Short than in Long task in all networks under hypotheses, e.g., for executive control (dACC) and valuation (vmPFC in Figure 3C).

**Table 2:** GLM1 overlapping clusters’ peak coordinates. Networks and areas were determined using probabilistic atlases (BA - Brodmann’s areas). No peak statistics was provided (see Method). Subclusters were omitted.

| Cluster ID | X | Y | Z | Network of Interest | Specific Area |
| --- | --- | --- | --- | --- | --- |
| Decision - Short & Long Overlap |  |  |  |  |  |
| 1 | 33 | -54 | 49.5 | Prospection | Right Precuneus |
| 2 | 63 | -39 | 23.25 |  | Right BA 22 |
| 3 | -48 | 30 | -10.5 | Executive Control | Left dIPFC |
| 4 | -6 | 54 | -14.25 | Executive Control | Left dACC |
| 5 | 27 | -39 | 64.5 | Prospection | Right Precuneus |
| 6 | -63 | -9 | -14.25 |  | Left BA 21 |
| 7 | 18 | -90 | -36.75 |  | Right Cerebellum |
| 8 | 27 | 51 | -14.25 | Executive Control | Right dIPFC |
| 9 | 60 | 0 | -18 |  | Right BA 21 |
| 10 | -21 | 42 | -18 | Prospection | Left dmPFC |
| 11 | -24 | -90 | -29.25 |  | Left Cerebellum |
| 12 | 48 | 39 | -14.25 |  | Right BA 47 |
| 13 | 6 | -33 | 60.75 |  | Right Superior Parietal Lobule |
| 14 | -66 | -27 | 0.75 |  | Left BA 21 |
| 15 | 54 | -27 | 4.5 |  | Right BA 22 |
| Confirmation - Short & Long Overlap |  |  |  |  |  |
| 1 | -42 | -24 | 60.75 |  | Left BA 4 |
| 2 | 0 | -54 | -33 |  | Midline Cerebellum |
| 3 | -6 | -24 | -6.75 |  | Left Thalamus |
| 4 | 30 | 27 | 4.5 |  | Right Insula |
| 5 | -30 | 27 | 0.75 |  | Left Insula |
| 6 | -21 | -93 | -10.5 |  | Left BA 18 |
| 7 | 12 | -84 | 0.75 |  | Right BA 17/18 |
| 8 | 42 | -72 | -21.75 |  | Right Cerebellum |
| 9 | -9 | -54 | 75.75 |  | Left BA 7 |
| 10 | -27 | -75 | -21.75 |  | Left Cerebellum |
| 11 | 45 | -51 | 60.75 |  | Right BA 40 |
| 12 | -15 | -87 | 42 |  | Left BA 19/7 |
| 13 | 6 | -45 | 19.5 | Prospection | Right PCC |
| 14 | -42 | -60 | 60.75 |  | Left Inferior Parietal Lobule |
| 15 | 33 | -3 | 53.25 |  | Right BA 6 |
| 16 | -15 | 21 | 15.75 | Executive Control | Left dACC |
| 17 | 21 | -66 | -40.5 |  | Right Cerebellum |
| 18 | -66 | -18 | 12 |  | Left Superior Temporal Gyrus |
| 19 | -57 | 21 | 12 |  | Left BA 45 |
| 20 | 24 | -42 | 64.5 |  | Right BA 5/7 |

During the confirmation epoch we found overlap in prospection (PCC) and executive control network (dACC) with more engagement of dlPFC and dACC, while less engagement of HC/MTL when Short was compared to Long task in Figure 3D. In addition, we looked at peak z-statistic within dACC on the individual subject level. Interestingly, the deactivation (absolute) peaks in decision were well correlated with activation peaks in confirmation across subjects (Pearson *r* = 0.58, *p <* 0.001).

No significant voxels survived with the two-sample t-test using GLM1 individual level BOLD response, giving no evidence towards hypothesized less overlap between short and long tasks for those subjects further away from the fitted line of log(*k_S_*) vs. log(*k_L_*).

The parametric modulation of reward revealed no significant clusters of activation during the decision epoch (GLM2), for the confirmation epoch in the Short task we found an increase in valuation network (vmPFC, Table 3) per unit increase in the reward value. Parametrically modulating delay magnitude we found negative modulation in dlPFC and positive modulation in dmPFC and vmPFC during the decision epoch for both tasks (Long in Figure 4A) with no significant clusters for confirmation. Finally, using *SV* for the parametric modulation we found that during the decision epoch the BOLD response amplitude in vmPFC was positively related to *SV* in the Short task, whereas ventral striatum activation was positively dependent on the *SV* in the Long task (Figure 4B). Under individual-level scrutiny, more subjects had peak z-statistic within valuation network in vmPFC than in ventral striatum for Short (17 out of 31), while for Long 17 subjects had peaks in ventral striatum and 11 subjects - in vmPFC, resulting in diverse group-level sensitivity across tasks.

**Table 3:** Significant clusters from parametric modulation: coordinates were provided with cluster size and peak statistics (only for non overlap, Method). Networks and areas were determined using probabilistic atlases (BA - Brodmann’s areas). Most subclusters were omitted.

| Cluster ID | X | Y | Z | Peak Stat | Cluster Size (mm<sup>3</sup>) | Network of Interest | Specific Area |
| --- | --- | --- | --- | --- | --- | --- | --- |
| GLM2c (Confirmation) Short |  |  |  |  |  |  |  |
| 1 | -48 | -66 | 4.5 | 6.9418 | 88931 |  | Left BA 19 |
| 2 | 66 | -33 | 19.5 | 6.8974 | 190721 |  | Right BA 22 |
| 3 | -12 | -75 | -51.75 | 5.2718 | 8842 |  | Left Cerebellum |
| 4 | -9 | 36 | -10.5 | 4.9210 | 7188 | Valuation | Left vmPFC |
| 5 | 0 | 9 | -6.75 | 4.9210 | 7188 |  | Left BA 25 |
| 6 | 57 | 33 | 4.5 | 4.5793 | 3881 | Executive Control | Right dlPFC |
| 7 | 30 | -33 | -18 | 4.4659 | 5096 |  | Right BA 37 |
| 8 | -54 | -6 | 45.75 | 4.3620 | 1586 |  | Left BA 6 |
| 9 | 21 | 0 | 53.25 | 4.3594 | 2362 |  | Right BA 6 |
| 10 | 15 | -72 | -55.5 | 4.3012 | 4117 |  | Right Cerebellum |
| 11 | 15 | -21 | 38.25 | 4.2283 | 1282 | Prospection | Right PCC |
| 12 | -15 | -33 | 38.25 | 4.1529 | 2430 | Prospection | Left PCC |
| 13 | -30 | 30 | -14.25 | 4.1058 | 1620 |  | Left BA 47 |
| GLM3 (Decision) Short & Long Overlap |  |  |  |  |  |  |  |
| 1 | -36 | -48 | 49.5 |  |  |  | Left BA 40 |
| 2 | 45 | -45 | 60.75 |  |  |  | Right BA 7 |
| 3 | 27 | -3 | 53.25 |  |  |  | Right BA 6 |
| 4 | -30 | -90 | 12 |  |  |  | Left BA 18 |
| 5 | -42 | 30 | 15.75 |  |  | Executive Control | Left dlPFC |
| 6 | -54 | 9 | 23.25 |  |  |  | Left BA 44 |
| 7 | 51 | 15 | 27 |  |  |  | Right BA 44 |
| 8 | -3 | 60 | 15.75 |  |  | Prospection | Left dmPFC |
| 8b | 0 | 54 | -6.75 |  |  | Valuation | Right vmPFC |

#### 3.2.1 ROI Analysis

Within the valuation ROI, we assessed individual beta-map correlations across all voxels between Short and Long tasks and averaged those across subjects (N = 30). One subject had an insignificant p-value and one subject had a negative correlation. Still, we found moderate positive correlation on average (*M* = 0.55, 95% CI [0.44, 0.65]).

Then, we tested whether *β*-coefficient averages from parametric modulation across peak voxels were significantly different from 0. For all models where peak voxels were found within the network (i.e. GLM2, GLM2c, GLM3, GLM4) the null hypothesis of *β*-coefficients being equal to 0 was rejected (all *ps <* 0.001) similarly for Short and Long tasks. We also established that averages of modulated parameter estimates for the peak voxels within the valuation network, such as reward for Short during confirmation, delay for Short during decision, and both Short and Long *SV* differed significantly between subjects’ groups closest to the fitted line of log(*k_S_*) vs. log(*k_L_*) and furthest (all *ps <* 0.001).

Individual differences analysis was also performed within the valuation ROI. We did not find any significant correlations between log(*k*) and modulated parameter estimates (all *ps >* 0.100). The only significant correlation we found was for beta-maps across tasks created by parametrically modulating the reward (Pearson *r* = 0.51, *p* = 0.003).

#### 3.2.2 RSA Analysis

Our whole-brain searchlight for representational similarity analysis (RSA) comparing neural RDMs to hypothesized dissimilarities in neural patterns between conditions (Figure 3B) identified eight brain clusters. These clusters showed similar activation pattern for Long and Short tasks during decision but differentiated between waiting and postponing, and included prospection network with HC/MTL (with coordinates (−60, −24, −22), (27, −27, −26), and (33, −6, −26)), dmPFC (−9, 42, 61), PCC/precuneus (−33, −42, 72), regions of cerebellum, and occipital cortex.

#### 3.2.3 Decoding Analysis

Decoding of | log(*k_Short_*) − log(*k_Long_*)| was equally well achieved using beta-maps from Short and Long tasks (Table 4). Cross-decoding, i.e., testing on the activation patterns from the opposite task, worked best when the model was trained on the beta-maps from the Short task across regions of interest. Overall, evaluating on the beta-maps from the other task yielded the highest performance within the executive control network explaining 71.5% of variance in absolute difference in discount factors.

**Table 4:** Decoding and cross-decoding *R*^2^ scores across brain networks.

| Region | Train → Test | $R^2$ |
| --- | --- | --- |
| Valuation | Short → Short | 0.979 |
| | Short → Long | 0.584 |
| | Long → Long | 0.979 |
| | Long → Short | 0.340 |
| Executive Control | Short → Short | 0.980 |
| | Short → Long | 0.715 |
| | Long → Long | 0.981 |
| | Long → Short | 0.553 |
| Prospection | Short → Short | 0.981 |
| | Short → Long | 0.521 |
| | Long → Long | 0.981 |
| | Long → Short | 0.377 |

Searchlight analysis over the joint mask of networks of interest predicting subjective value (*SV*) with trial-by-trial beta-maps was reported in Figure 5A. The quality of decoding as measured by R² was poor explaining less than 4% of variance.

**Figure 5:**
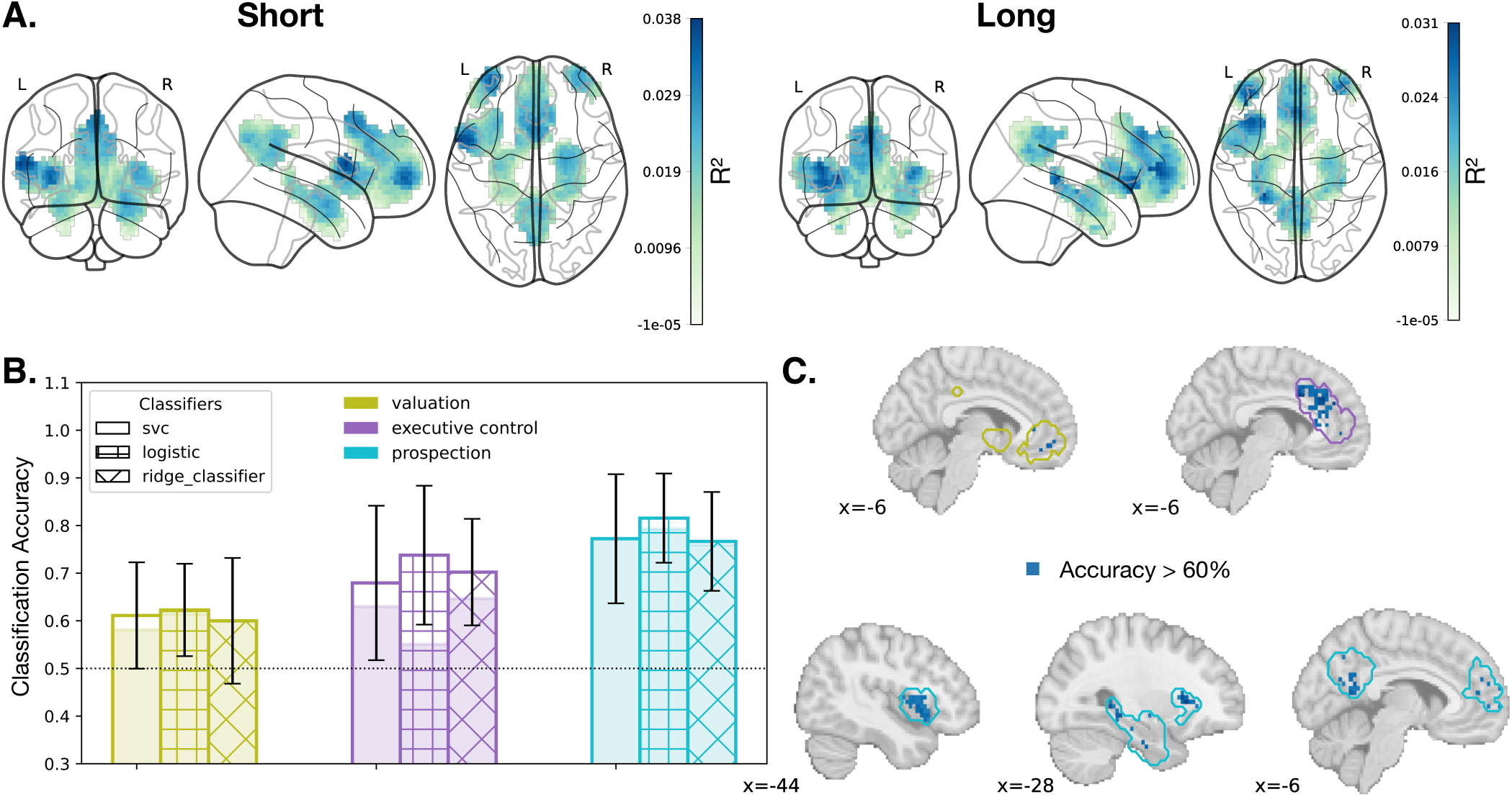
Decoding results. **A.** Prediction performance maps showing voxel-wise R² scores from support vector regression (SVR)-based searchlight analysis aggregated across participants. Trial-wise beta estimates were used to predict subjective value (SV) in the Short (left) and Long (right) delay tasks. **B.** Training (bar height) and testing (shaded area) CV-accuracy for classifying Short vs Long tasks using three models and beta maps covering three networks: valuation, executive control, and prospection. Pattern for the bar defined the classifier. The chance level was given by a dotted horizontal line. Classification within prospection ROI achieved the highest testing accuracy of 79%. **C.** Accuracy maps from the searchlight analysis thresholded at 60% (ROC-AUC). Color contour identified the brain network (same as in B). vmPFC was highlighted within the valuation ROI (at *x* = −6, top-left). dACC was identified as a key area for executive control (also at *x* = −6, top-right). Within the prospection network all areas contributed to the classifier’s prediction (brain images at x = [−44, −28, −6], bottom row).

Classification of the tasks worked best within the prospection network achieving 82% training accuracy and 79% testing accuracy (with logistic regression being the best), whereas executive network beta-maps achieved 65% testing accuracy and valuation network got a maximum of 61% testing accuracy (Figure 5B). The searchlight analyses achieved smaller accuracies partially due to distributed (rather than localized) signals with prospection network being the best and reaching 78% accuracy. In prospection network the accuracy was driven cumulatively by all areas in the network, while the valuation network highlighted the role of vmPFC driving the model accuracy and the executive control - dACC (Figure 5C).

## 4 Discussion

In our study we aimed to explore the neural correlates of intertemporal choices involving waiting and postponing. Behavioral results were consistent with our previous research. The discount factors across tasks were correlated, but not related to self-reported impulsivity (BIS) or the decision response times. After eliminating subjects at extreme, log(*k_S_*) and log(*k_L_*) in the fMRI study were no longer significantly correlated. Still, the individual discount factors between computer and fMRI sessions were stable.

Statistical modeling of imaging data partially supported our first hypothesis by finding extensive activation overlap in executive control and prospection networks across tasks during decision making and waiting. We further supported this hypothesis with the valuation network ROI analysis, showing similar pattern of activation across tasks via significant positive correlations of individual beta-maps from nonmodulated epoch regressors and tests of significance of *β*-coefficients for parametrically modulated regressors. Contrary to our hypothesis, we found significant differences across tasks within the valuation network (the key circuit for the integration and calculation of subjective value as in Hare et al., 2008, 2014; Lukinova & Myagkov, 2016). Importantly, there was less activation in vmPFC for Short compared to Long task during choice. We attributed that to the epoch-related GLM analysis in our study compared to the choice-related one highlighting the vmPFC role (e.g., Jimura et al., 2013).

We did not find support for our second hypothesis in the whole-brain analysis. There were no significant clusters that differentiated between subjects closer and those further away from the fitted line of two discount factors. In the ROI analysis, we found consistent significant differences between average modulated parameters only for the Short task for the groups of subjects as above, suggesting a subtle support for the hypothesis that would need to be confirmed in bigger cohorts.

With our parametric modulation analyses we supported hypothesis three, in particular, following Kable and Glimcher (2007) we concluded that valuation network was positively modulated by subjective value at the time of choice, highlighting the role of vmPFC and ventral striatum. Interestingly, we found amplitude of vmPFC activation being positively related to the reward for waiting and payment period, probably suggesting the coins salience right after waiting.

Our a priori-defined analyses regarding correlations between neural activation and individual discounting parameters testing the fourth hypothesis did not yield any significant findings. This could be explained by the exclusion of subjects with extreme discount factors before fMRI session. Previous studies were successful in registering a small correlation of *r* ∼ .40 for a much larger cohort (*N >* 100, Koban et al., 2023). Within the valuation ROI mask we were only able to register a significant correlation between parametric *β*-weights for reward across tasks and not for other parametric modulations, suggesting that the neural encoding of delays and subjective values of delayed rewards captured by parametric modulation was diverse across temporal contexts.

We also had several additional weaker hypotheses, mostly focused on the networks of executive control and prospection. As anticipated we found negative parametric modulation of dlPFC (as in Ballard & Knutson, 2009; Wang et al., 2021, hypothesis A1) and positive parametric modulation of dmPFC (prospection network, A2) by delay in both tasks. We also supported hypotheses about stronger engagement of executive control network (A3), but smaller activations in the prospection region (A4) for Short compared to Long tasks after choice during the confirmation epoch that included waiting.

Although we did not find strong significant correlations between neural activity and discount factors, this would not necessarily mean no overlap in neural underpinnings. Likewise, finding a significant overlap in brain regions engaged in both Short and Long tasks might not be a strong evidence for a common code. With multi-voxel pattern analysis, we checked whether the similarities and differences across those neural representations were distinct at a more fine-grained level. We found prospection network being crucial both to temporal proximity of the tasks and their divergence.

The prospection network is central to the concept of ‘mental time travel’, allowing individuals to simulate future outcomes and integrate episodic memory into intertemporal choices (Boyer, 2008). Prior research demonstrated that episodic future thinking could reduce temporal discounting (Benoit et al., 2011; Peters & Büchel, 2010), suggesting that activation in this network might underlie both the shared and distinct neural patterns observed across time scales. While valuation processes are crucial for assigning subjective worth to future rewards (Lempert et al., 2018), the prospection network enables individuals to bridge temporal gaps, promoting long-term decision-making (Lempert, n.d.).

This study was the first to look at intertemporal choice task across short and long delays. We compared tasks, not considering brain activity related to choices. Further studies need to establish whether the engagement of the prospection network explains differentiated recruitment depending on the cognitive demands of future-oriented decisions and the precise functions of the regions within the network in the formation of time preferences under usual or very short delays.

## Data and Code Availability

The tasks were coded in PsychoPy toolbox (3.1.5, Peirce, 2007) and run on a Windows 7 laptop. All analyses, statistics, and visualizations were performed either in Matlab (version 24.2, or higher, The Mathworks, MA), in R (version 4.4.0, or higher, R Foundation for Statistical Computing, Vienna, Austria), or in Python (version 3.12.7). The fMRI data was preprocessed using fMRIPrep (version 23.2.3) and analyzed using Nilearn (version 0.10.4). Code and behavioral data are available through a GitHub repository (https://www.github.com/evgluk/fMRIPvsW).

## Author Contributions

All authors contributed to the study design and preregistration. Data collection and analyses were performed by SX and EL. EL wrote the manuscript. All authors reviewed the manuscript.

## Acknowledgements

We acknowledge outstanding undergraduate students who helped us to start the project and to collect the data: Danielle John from CUNY Hunter College, Brianna Fu & Yui Ying (Chloe) Wong from New York University, and Xirui Zhang, Xue Bai & Weiyi (Molly) He from New York University Shanghai.

